# A red light-induced genetic system for control of extracellular electron transfer

**DOI:** 10.1101/2023.12.02.569691

**Authors:** Fengjie Zhao, Christina M. Niman, Ghazaleh Ostovar, Marko S. Chavez, Joshua T. Atkinson, Benjamin M. Bonis, Jeffrey A. Gralnick, Mohamed Y. El-Naggar, James Q. Boedicker

## Abstract

Optogenetics is a powerful tool for spatiotemporal control of gene expression. Several light-inducible gene regulators have been developed to function in bacteria, and these regulatory circuits have been ported into new host strains. Here, we developed and adapted a red light-inducible transcription factor for *Shewanella oneidensis*. This regulatory circuit is based on the iLight optogenetic system, which controls gene expression using red light. Promoter engineering and a thermodynamic model were used to adapt this system to achieve differential gene expression in light and dark conditions within a *S. oneidensis* host strain. We further improved the iLight optogenetic system by adding a repressor to invert the genetic circuit and activate gene expression under red light illumination. The inverted iLight genetic circuit was used to control extracellular electron transfer (EET) within *S. oneidensis*. The ability to use both red and blue light-induced optogenetic circuits simultaneously was demonstrated. Our work expands the synthetic biology toolbox of *Shewanella*, which could facilitate future advances in applications with electrogenic bacteria.

## Introduction

Optogenetics combines light-sensitive proteins and genetic techniques to control cellular processes within living organisms^1^. Synthetic optogenetic circuits can be constructed to tune gene expression with light-responsive transcription factors^1^. In recent years, optogenetic circuits have been developed to control gene expression in bacteria in response to illumination with blue, green, red, or near-infrared light^2–7^. Optogenetic circuits have been utilized to control gene expression to regulate many microbial processes, such as biochemical production^8–11^, biofilm formation^12,13^, and bacterial infection^14^. Although optogenetics has been implemented in model bacteria, such as *Escherichia coli*^6,12^, *Bacillus subtilis*^15^ and *Pseudomonas aeruginosa*^14,16^, there is still a need to extend these systems to other hosts for specific microbial processes.

*Shewanella oneidensis* MR-1 is a model electroactive organism, whose extracellular electron transfer (EET) pathways have been well studied^17,18^. EET within *S. oneidensis* MR-1 utilizes a network of multiheme *c*-type cytochromes to route electrons from the cellular interior to external electron acceptors^19–24^. Synthetic biology strategies have recently been developed to regulate the EET capabilities of *S. oneidensis*, such as using genetic circuits to control the genes encoding the multiheme *c*-type cytochromes of the EET pathway. West *et al*., developed a native inducible system to tune the expression level of one multiheme cytochrome-porin complex (MtrCAB) to control EET capabilities^25^. In another study, clustered regularly interspaced short palindromic repeats interference (CRISPRi) and small regulatory RNA (sRNA) were used to repress the transcription and translation of *mtrA* to regulate EET efficiency^26^. Recently, Dundas *et al*., developed chemical induced transcriptional logic gates to control the EET flux of *S. oneidensis* through tuning the transcription and translation of EET related genes^27^. A plasmid toolkit with different promoters and replication origins were characterized and utilized to control cytochromes expression in *S. oneidensis* for EET regulation^28^. The EET pathway of *S. oneidensis* MR-1 had also been reconstructed successfully in *E. coli* through heterologous expression of the related genes encoding the *c*-type cytochromes^29,30^. Despite these advances, synthetic biology toolboxes, in particular the use of optogenetic gene circuits to control EET, are still limited in *S. oneidensis*.

*S. oneidensis* can transfer electrons through cytochromes on the nanometer-scale and can also form living conductive biofilms for long-distance electron transport across neighboring cells on the micrometer-scale^31–33^. We previously developed a lithographic strategy to pattern conductive biofilms of *S. oneidensis*, using the blue light-induced genetic circuit pDawn to control cell aggregation^34^. This technique enabled tunable current generation by varying the dimensions of electroactive biofilms. Since our previous work showed the potential of using optogenetics to pattern electrogenic microbes on the micrometer scale, adapting additional optogenetic systems for *S. oneidensis* could enable new strategies to control EET and develop advanced living electronics.

A recently published paper showed a single-component red light-induced optogenetic system, iLight, for transcriptional regulation in *E. coli*^35^. We developed and adapted this iLight optogenetic system into *S. oneidensis* through promoter engineering and a thermodynamic model. Then we improved the iLight optogenetic system to activate the gene expression using red light in *S. oneidensis*. Finally, we used this iLight genetic circuit to control the expression of cytochromes and regulate the EET activity of *S. oneidensis*. Our work demonstrated a new circuit to control gene expression in *S. oneidensis* using red light and demonstrated light-induced EET activity of *S. oneidensis*.

## Results

### Importing the iLight optogenetic system to *S. oneidensis*

The iLight optogenetic system was originally developed and optimized in *E. coli*^35^. It consists of a plasmid that encodes a light-sensitive repressor. The repressor contains a LexA408 DNA-binding domain fused to a photosensory domain IsPadC-PCM. This protein has a sfGFP fused to the C-terminus. An RFP reporter gene was regulated by the iLight repressor (Fig. 1A). The proposed mechanism of action of the iLight optogenetic system was red light induced tetramerization of iLight photosensory module, which enabled the LexA408 DNA-binding domains to form active transcription factor dimers^35^. The iLight photosensory module requires biliverdin IXα (BV) tetrapyrrole as the chromophore to enable the light-responsive function of the iLight repressor (Fig. S1). In our system, the *ho1* gene, a heme oxygenase, is expressed from the iLight plasmid for BV synthesis (Fig. 1A).

**Figure 1.**
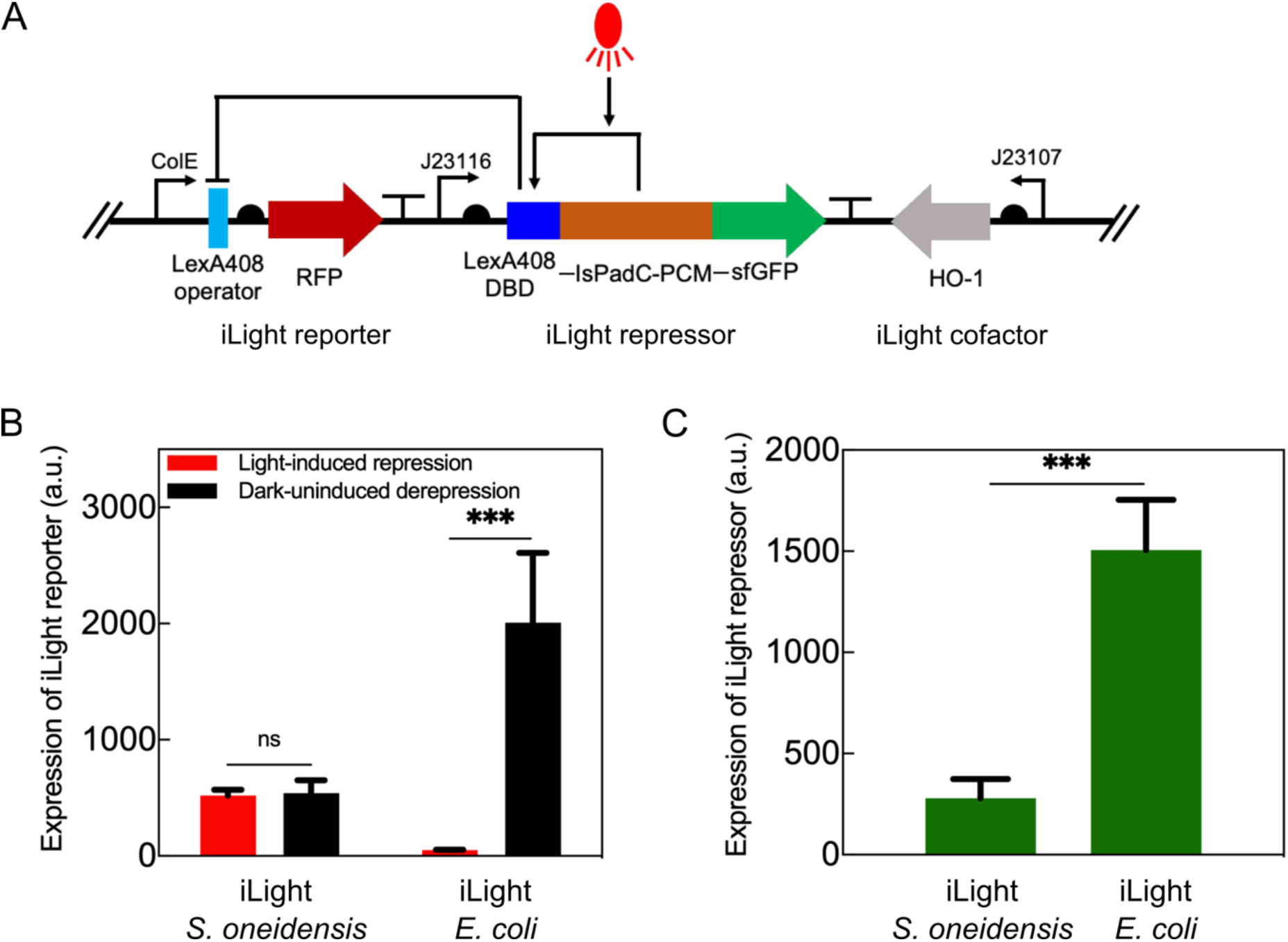
Characterization of the iLight genetic circuit in *S. oneidensis* and *E. coli*. (A) The single plasmid iLight genetic circuit contained iLight repressor, iLight reporter and iLight cofactor. The RFP reporter measures light-regulated gene expression and sfGFP measures the expression level of the iLight repressor. (B) RFP fluorescence intensity measurements of iLight reporter expression in *S. oneidensis* and *E. coli* cultured under red light and dark conditions, respectively. *p* = 0.7462 for *S. oneidensis* Light vs *S. oneidensis* Dark and *p* = 0.0006 for *E. coli* Light vs *E. coli* Dark (two-tailed unpaired *t* test). (C) GFP fluorescence intensity measurements of iLight repressor expression in *S. oneidensis* and *E. coli. P* = 0.0005 for *S. oneidensis* vs *E. coli* (two-tailed unpaired *t* test). The measurements (mean ± SD) were derived from triplicate experiments. a.u., arbitrary units. Significance is indicated as \*\*\**p* < 0.001 and ns (not significant) *p* > 0.05.

We then tested this single plasmid iLight genetic circuit in *E. coli* and *S. oneidensis*. In *E. coli*, RFP fluorescence measurements confirmed the iLight genetic circuit repressed expression of the RFP reporter gene under illumination of red light (Fig. 1B). When cultured in the dark, the RFP expression was derepressed resulting in 40-fold increase in RFP expression (Fig. 1B). However, in *S. oneidensis* transformed with the iLight plasmid, the expression level of the RFP reporter was similar in both cells cultured in red light and in the dark (Fig. 1B). To determine if this circuit failure was caused by limited expression of the iLight repressor in a new host, we measured expression using the sfGFP fusion to iLight. The expression level of iLight repressor in *S. oneidensis* was much lower than in *E. coli* (Fig. 1C). We hypothesized that the low expression of the iLight repressor could be the reason why the iLight genetic circuit did not result in light-regulated gene expression in *S. oneidensis*.

### Adjusting the expression level of the iLight repressor

The expression of iLight repressor was much lower in *S. oneidensis* than in *E. coli*, suggesting differential gene expression in light and dark conditions might depend on the level of repressor expression. To test this hypothesis, we first developed a thermodynamic model to predict how the expression level of the iLight repressor impacts the fold change of the iLight reporter when exposed to dark and light conditions (see Supplementary Methods and Fig. S2). The thermodynamic model^36-38^ investigates RFP gene expression considering the binding probabilities of RNA polymerase and iLight repressor molecules to the promoter region in red light and dark conditions (Fig. S2A). Additionally, it assumes that the dimeric form of the iLight repressor in dark exhibits a non-zero, but lower, probability of binding to the specific site compared to tetrameric form in red light. We found that the fold change in gene expression was largest for intermediate expression levels of iLight repressor (Fig. S2B and C), suggesting that increasing expression of the iLight repressor might improve the performance of this optogenetic circuit within *S. oneidensis*.

The expression of the iLight repressor was modified by site directed mutagenesis of the promoter region (Fig. 2A). The original promoter J23116 was weak^39^. Based on the Anderson promoter collection (http://parts.igem.org/Promoters/Catalog/Anderson), the promoter of the iLight repressor was mutated to achieve a broad range of expression levels. As shown in Fig. 2B, the modified promoters varied in expression of iLight repressor over 44-fold. Expression of the RFP iLight reporter gene in both light and dark conditions was measured for each of these promoters (Fig. S3). Promoters J23102 and J23108, which expressed intermediate levels of iLight repressor, showed the largest fold change of the iLight reporter (Fig. 2C). The fold changes of iLight reporter (dark/red light) were around 12 for both iLight-J23102 and iLight-J23108 (Fig. 2C). We also tested the performance of these iLight genetic circuits with modified expression of the iLight repressor in *E. coli*. The fold change in expression of the iLight reporter decreased to 1 at high levels of iLight repressor expression (Fig. S4). Unlike in *S. oneidensis*, the weakest promoters resulted in fold changes above 20, likely due to these weak promoters having higher expression in *E. coli* than in *S. oneidensis*. In summary, adaption of the iLight optogenetic system for use in *S. oneidensis* required adjustment of the expression level of the iLight repressor.

**Figure 2.**
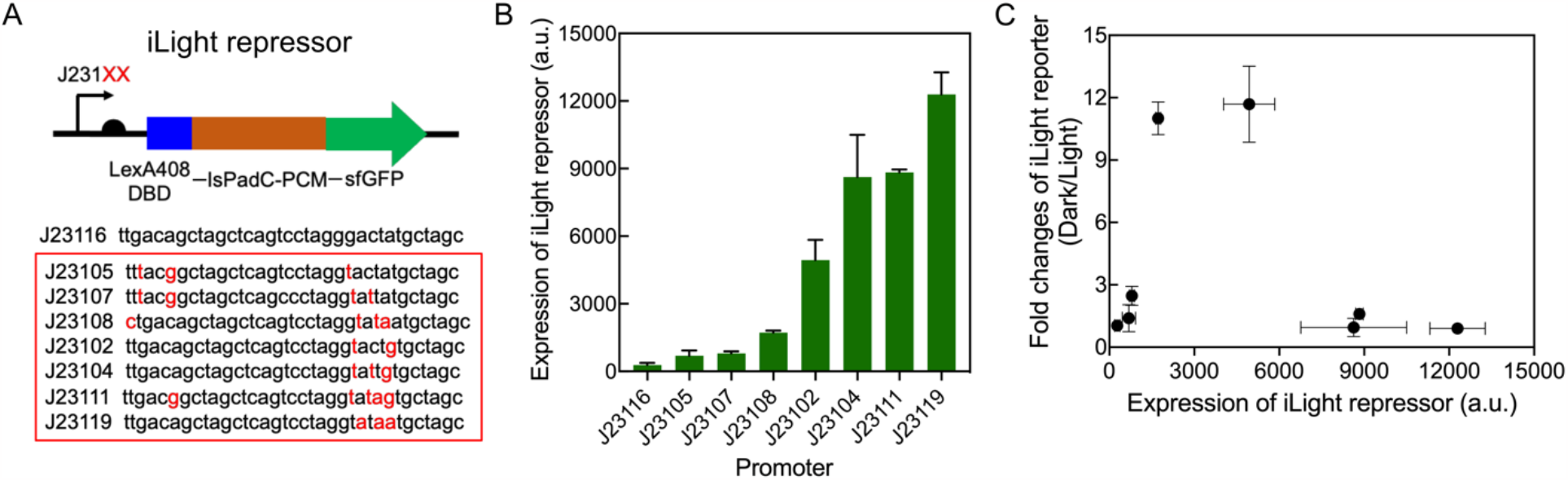
Adapting the iLight optogenetic system for *S. oneidensis*. (A) Site directed mutagenesis of the promoter to tune expression of the iLight repressor. (B) Expression of the iLight repressor in *S. oneidensis* from these promoters, as measured via expression of co-transcribed sfGFP. (C) Fold changes of the iLight reporter (dark/red light) in *S. oneidensis* strains containing different promoters of iLight repressor. The measurements (mean ± SD) were derived from triplicate experiments. a.u., arbitrary units.

### Creating an inverted iLight optogenetic system, for light-activated gene regulation

The iLight optogenetic system could work in *S. oneidensis* MR-1 after adjusting the expression level of the iLight repressor. In the original report, the iLight system was used to repress gene expression in *E. coli* with red light (Fig. 3A), but other optogenetic circuits have been modified for both activation and repression with light inputs. Next, we improved the iLight optogenetic system to activate the target gene expression through red light illumination in *S. oneidensis* MR-1.

**Figure 3.**
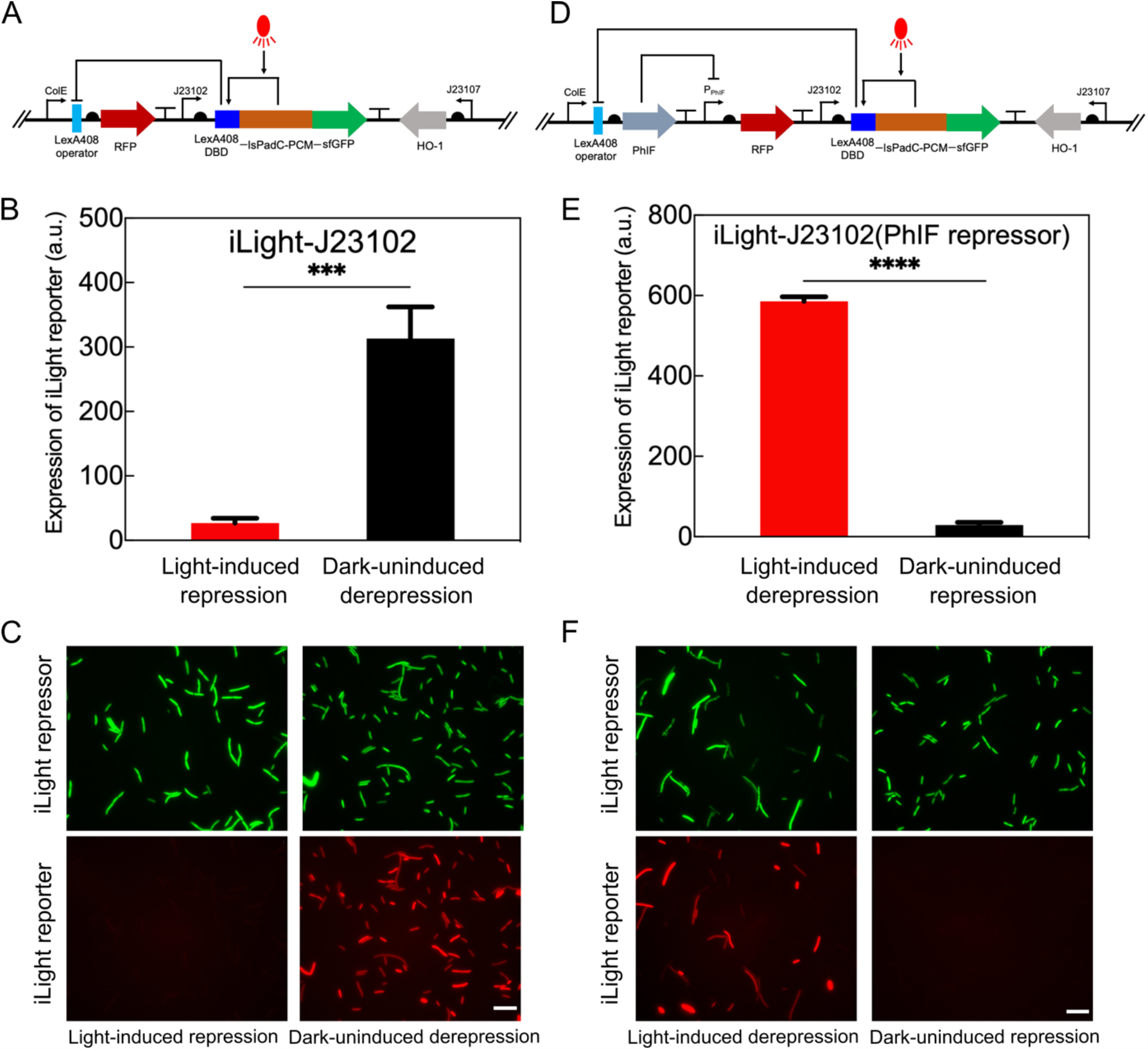
Characterization of the non-inverted iLight genetic circuit iLight-J23102 and the inverted iLight genetic circuit iLight-J23102(PhIF repressor) in *S. oneidensis*. (A) Genetic circuit of non-inverted iLight-J23102. (B) RFP fluorescence intensity measurements of iLight reporter expression in the non-inverted iLight-J23102 strain cultured under red light and dark conditions, respectively. *p* = 0.0006 for Light vs Dark (two-tailed unpaired *t* test). (C) Microscope observation of iLight reporter and iLight repressor for non-inverted iLight-J23102 cells cultured under red light and dark conditions, respectively. Scale bars 10 μm. (D) Genetic circuit of inverted iLight-J23102(PhIF repressor). (E) RFP fluorescence intensity measurements of iLight reporter expression in the inverted iLight-J23102(PhIF repressor) strain cultured under red light and dark conditions, respectively. *p* < 0.0001 for Light vs Dark (two-tailed unpaired *t* test). (F) Microscope observation of iLight reporter and iLight repressor for inverted iLight-J23102(PhIF repressor) cells cultured under red light and dark conditions, respectively. Scale bars 10 μm. The measurements (mean ± SD) were derived from triplicate experiments. a.u., arbitrary units. Significance is indicated as \*\*\**p* < 0.001 and \*\*\*\**p* < 0.0001.

To accomplish this, we incorporated the iLight repressor into an inverter genetic circuit by adding a second repressor. Expression of the second repressor regulates the target gene and is regulated by iLight repressor. After red light illumination, the expression of this second repressor is repressed by the iLight repressor, resulting in derepression of the target gene (Fig. 3D). We tested three different repressors, the λ phage repressor cI and the TetR-family repressors PhIF and SrpR^40^, with their cognate promoters to invert the iLight-J23102 genetic circuit. All three repressors resulted in increased expression of the target gene in response to red light (Fig. 3E and Fig. S5A-C). The fold changes of iLight reporter RFP between red light and dark conditions were 16 for cI repressor, 20 for PhIF repressor and 12 for SrpR repressor (Fig. 3E and Fig. S5A-C). Microscopic images showed that the non-inverted iLight-J23102 cells had strong red fluorescence after being cultured under dark but weak fluorescence after being cultured with red light (Fig. 3C), as expected for light-induced repression of gene expression. The microscopic images of inverted iLight-J23102(PhIF repressor) strain showed the cells had strong red fluorescence after being cultured with red light, but weak fluorescence after being cultured in the dark (Fig. 3F), as expected for light-induced activation of gene expression. We also found that inverting the iLight-J23108 genetic circuit using the cI repressor resulted in just 5-fold change in expression of the reporter gene (Fig. S5D), which was much lower than those of inverting iLight-J23102 (Fig. S5A). The reason for this could be higher leaky expression of the inverting repressor (cI) for iLight-J23108 relative to iLight-J23102 under red light condition (Fig. S3). We selected the iLight-J23102(PhIF repressor) genetic circuit for subsequent experiments, which had the highest fold change.

We then checked whether *S. oneidensis* could utilize two different light-regulated genetic constructs responding to different colors of light simultaneously. To achieve this, we introduced the blue light-induced pDawn^3^ genetic circuit to control GFP expression and red light-induced iLight genetic circuit to control RFP expression within *S. oneidensis*. The iLight circuit used here did not have the sfGFP fusion to the iLight repressor. As we expected, *S. oneidensis* cells containing both iLight and pDawn genetic circuits had strong red fluorescence only when cultured with red light and strong green fluorescence only when cultured with blue light (Fig. S6).

Taken collectively, we inverted the iLight optogenetic system to activate the target gene expression under red light illumination in *S. oneidensis* and the fold change was increased compared with the non-inverted iLight genetic circuit. The inverted iLight genetic circuit could work together with blue light-induced pDawn genetic circuit to control the expression of two different genes with two lights, respectively.

### The extracellular electron transfer activity of *S. oneidensis* can be regulated with red light

*S. oneidensis* MR-1 can transport electrons from cytosolic metabolism to external electrodes using an EET pathway involving *c*-type cytochromes located in the inner membrane, periplasm, and outer membrane^18^ (Fig. 4A). To determine if we can control the EET activity of *S. oneidensis* using light, we used the iLight optogenetic system to control the expression of multiple cytochromes within this EET pathway. We constructed *S. oneidensis* strain iLight-MtrC which utilized the inverted iLight-J23102(PhIF repressor) genetic circuit to activate the expression of outer membrane cytochrome MtrC with red light. The host strain for this construct was *S. oneidensis* Δ*mtrC*Δ*omcA*^41^ in which the key genes encoding outer membrane cytochromes were deleted from the genome. MtrC expression from the promoter was evaluated by measuring the fluorescence due to the RFP gene co-transcribed from the same promoter (Fig. S7A). As expected, the RFP fluorescence was much higher when culturing iLight-MtrC strain under red light illumination, while the RFP fluorescence was weak when culturing the same strain under dark condition (Fig. S7B). These results suggest that MtrC could be expressed under red light illumination with very low expression in the dark.

**Figure 4.**
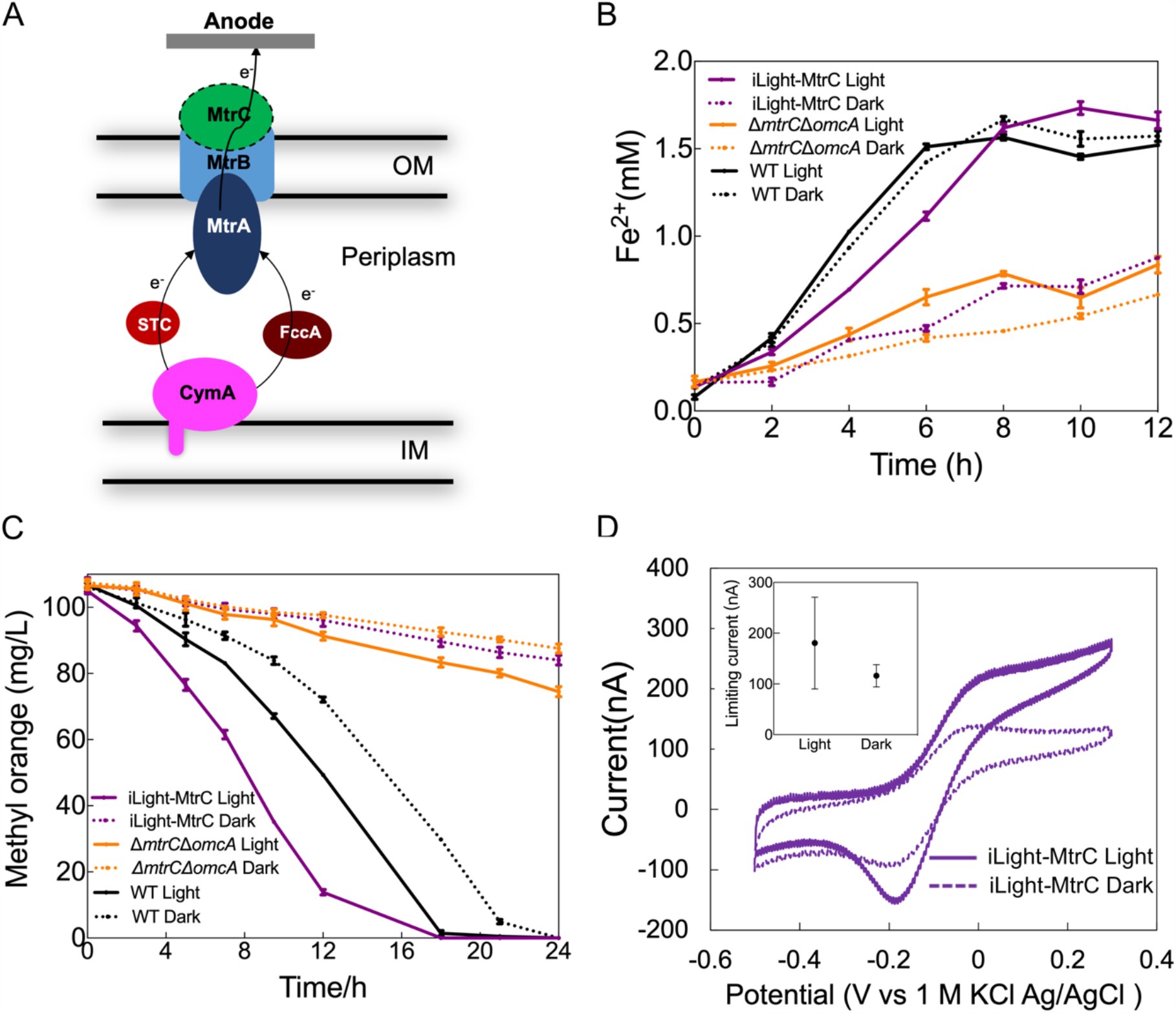
Using iLight-J23102(PhIF repressor) genetic circuit to control the outer membrane cytochrome MtrC expression for light-induced extracellular electron transfer (EET) activity in *S. oneidensis*. (A) The EET pathway of *S. oneidensis* MR-1. (B) Iron reduction assay for iLight-MtrC strain, Δ*mtrC*Δ*omcA* and wild type with blank plasmid iLight-J23102(PhIF repressor) after being cultured under red light and dark conditions, respectively. (C) Methyl orange decoloration assay for iLight-MtrC strain, Δ*mtrC*Δ*omcA* and wild type with blank plasmid iLight-J23102(PhIF repressor) after being cultured under red light and dark conditions, respectively. (D) Cyclic voltammetry curves for iLight-MtrC strain after being cultured under red light and dark conditions, respectively. The inset graph showed the average limiting current from cyclic voltammetry measurements of three biological replicates. Data shows mean ± SD from triplicate experiments.

Next, EET activity was measured for the iLight-MtrC strain cultured under light and dark conditions using colorimetric assays for extracellular redox activity. *S. oneidensis* can perform anaerobic respiration by reducing a wide range of external terminal electron acceptors^42–47^. We utilized iron citrate and methyl orange (MO) as the electron acceptors to measure the Fe^3+^ and azo dye reduction capabilities of the iLight-MtrC strain. As shown in Fig. 4B and C, the iLight-MtrC strain had much higher Fe^3+^ and MO reduction rates when cultured under red light than those of iLight-MtrC grown in the dark. The reduction activities of red light illuminated iLight-MtrC cells were comparable with those of wild type cells cultured either under red light or in the dark. The reduction activities of iLight-MtrC cells grown in the dark were as low as those of Δ*mtrC*Δ*omcA* cultured either under red light or in the dark (Fig. 4B and C). These results indicated the reduction activities of the iLight-MtrC strain could be controlled by light.

Finally, we performed electrochemical measurements to determine if light-induced expression of MtrC would modulate current production in *S. oneidensis* biofilms. Cells were grown in bioreactors with planar indium tin oxide (ITO) coated glass coverslips as bottom with or without red light illumination. Cyclic voltammetry (CV) was performed for the biofilms on the ITO electrodes. As shown in the cyclic voltammetry curves in Fig. 4D, we found higher current for red light illuminated iLight-MtrC cells than the same strain grown in the dark. The average limiting current from cyclic voltammetry measurements of three biological replicates were 180 nA for red light illuminated cells and 110 nA for cells cultured under dark (Fig. 4D inset graph).

We also constructed *S. oneidensis* strain iLight-STC to use the iLight-J23102(PhIF repressor) genetic circuit to control expression of the periplasmic small tetraheme cytochrome (STC, encoded by the gene *cctA*)^48^ by red light within *S. oneidensis* Δ*stc*Δ*fccA*. STC could be expressed under red light illumination with very low expression in the dark (Fig. S8A). The Fe^3+^ reduction activities of the iLight-STC strain could also be controlled by red light (Fig. S8B). These results demonstrate that we can regulate the EET activity of *S. oneidensis* using optogenetic circuits that regulate cytochrome expression.

## Discussion

We developed and adapted a red light-induced genetic circuit for *S. oneidensis* MR-1 based on a previously reported iLight optogenetic system^35^. The iLight genetic circuit reported for *E. coli* did not initially work in *S. oneidensis*, with no fold-change of gene expression in response to light. Through modeling and experimental tests, we found that the iLight genetic circuits required a specific expression range of the light-responsive transcription factor to function properly. Our modeling suggested that an intermediate level of the repressor would be needed for light-regulated gene expression. Low expression of iLight repressor could result in insufficient formation and binding of the repressor tetramer under the red light condition, which led to insufficient repression of the iLight reporter. High expression of iLight repressor resulted in repression of the regulated gene under both dark and light conditions, as the large concentration of the repressor compensated for the weak binding of the repressor in the dark state. Tuning of the expression level of the iLight repressor via promoter engineering identified constructs with differential expression of the reporter gene in light and dark conditions. This work demonstrated how the expression levels of transcription factor proteins can be critical to the function of a genetic circuit, and similar promoter optimization may be needed to adapt the iLight optogenetic system in other bacterial host strains.

To expand the regulatory capability of the iLight system in *S. oneidensis*, we inverted the iLight genetic circuit by adding an additional repressor to activate the gene expression by red light (Fig. 3). We selected three different repressors with their cognate promoters^40^. Inverting iLight with cI and PhIF repressors achieved higher expression of RFP than with a circuit using the SrpR repressor. The fold changes in gene expression were greater in the inverted iLight genetic circuit containing cI and PhIF repressors than in the non-inverted iLight. The reason could be the cognate promoters of cI and PhIF repressors are stronger than the ColE promoter, so expression levels of RFP are higher in inverted iLight under red light than that of non-inverted iLight under dark. A similar trend was observed for the blue light-induced pDawn and pDusk genetic circuits^3^.

Prior efforts to regulate the EET activity of *S. oneidensis* usually depended on using chemically induced genetic circuits to control the expression of cytochromes^25,27,28^. Compared with chemical induced genetic circuits, light-induced genetic circuits make it possible to create spatial patterns of microbial activity^9,10,12,13^. Our work using the red light-induced iLight genetic circuit to control cytochromes expression further expands the synthetic biology toolboxes of *S. oneidensis* and demonstrates light-induced EET activities, which can be a promising approach to spatiotemporally control the EET of *S. oneidensis*. This work allowed us to tune the electron transfer using light. Developing optogenetic tools for *S. oneidensis* will have implications for both studying and harnessing bioelectronics and the development of advanced living electronics.

## Methods

### Bacterial strains and plasmids

*Escherichia coli* DH5α and NEB stable were used for plasmid construction. *Shewanella oneidensis* MR-1 was used as the host to characterize the performances of different non-inverted and inverted iLight constructs in *S. oneidensis. S. oneidensis* Δ*mtrC*Δ*omcA* (strain JG749)^41^ was used as the host to contain the plasmid with expression of outer membrane cytochrome MtrC controlled by the inverted iLight genetic circuit. *S. oneidensis* Δ*stc*Δ*fccA* (strain JG3107) was used as the host to contain the plasmid with expression of periplasm cytochrome STC controlled by the inverted iLight genetic circuit. The Δ*stc*Δ*fccA* strain was generated by deleting *fccA* using materials and methods described previously^49^ in strain JG561, a background where *cctA* has been deleted^41^. *E. coli* NEB stable was used as the host to characterize the performances of different non-inverted iLight constructs in *E. coli*.

The original iLight plasmid^35^ was purchased from Addgene (Catalog #170268). A *ho1* gene was amplified from plasmid pNO41^5^ (Addgene, Catalog #101067) and then added to the original iLight plasmid to create iLight-J23116 plasmid. To construct iLight plasmids with different promoters transcribing the iLight repressor, iLight-J23116 was used as the starting plasmid. Site directed mutagenesis was then performed for the promoter region of iLight repressor through using NEB Q5® Site-Directed Mutagenesis Kit with non-overlapping primers or NEBuilder® HiFi DNA Assembly (New England BioLabs, MA, USA) with overlapping primers. cI, PhIF and SrpR repressors with their cognate promoters were amplified from plasmid pDawn-mCherry^34^, pRF-PhIF^40^ (Addgene, Catalog #49367) and pRF-SrpR^40^ (Addgene, Catalog #49372), respectively. Then, these DNA fragments were added to the iLight-J23102 and iLight-J23108 plasmids after the LexA408 operator to obtain different inverted iLight genetic circuits. To test whether *S. oneidensis* could utilize iLight and pDawn genetic circuits to respond to red light and blue light simultaneously, we constructed a plasmid to control GFP expression by pDawn. We removed the GFP reporter for the expression of the iLight repressor and changed the antibiotic resistance from spectinomycin to kanamycin in the iLight plasmid. For constructing light-induced EET plasmids, *mtrC* and *stc* genes were amplified from the genome of *S. oneidensis* MR-1 and added before the *mCherry* gene in the inverted iLight plasmids under the control of the cognate promoter of the second repressor.

All strains, plasmids and primers used in this study are listed in Table S1, S2 and S3.

### Growth conditions

*E. coli* strains were cultivated in Lysogeny broth (LB) medium (10 g/L tryptone, 5 g/L yeast extract and 5 g/L sodium chloride) at 37 °C, 200 rpm. *S. oneidensis* strains were cultivated in LB medium or minimal medium at 30 °C, 200 rpm. The minimal medium recipe can be found in Table S4. When necessary, media were supplemented with spectinomycin (Spec, 100 μg/mL) and kanamycin (Kan, 50 ug/mL).

For anaerobic culturing of *S. oneidensis*, the minimal media with different kinds of electron acceptors were purged with nitrogen. The minimal medium without vitamin solution was purged with nitrogen and used for the electrochemical measurements. The anaerobic culturing was performed in the sealed serum bottles and electrochemical measurements were performed in an anaerobic chamber.

### Fluorescence measurements and microscopy

To characterize the performance of iLight genetic circuits in *E. coli* and *S. oneidensis*, overnight cultures (1%, v/v) of the strains were transferred into 5 mL fresh LB broth and grown to late log phase (OD^600nm^ about 1-1.5). Then, the cultures (1%, v/v) were seeded into 5 mL of LB broth and incubated at 37 °C or 30 °C, for *E. coli* and *S. oneidensis*, respectively, either under the red light or dark condition while shaking at 200 rpm. Red light was provided by attaching LED strip lights to the wall inside the shaker with an intensity of 150 μW/cm^2 35^. *S. oneidensis* containing both the pDawn and iLight genetic circuits were exposed to blue light and red light, both with intensities of 150 μW/cm^2^. An optical power meter (PM100USB, Thorlabs) was used for measuring the illumination intensity of lights. Cultures were collected after 18 h incubation to measure the fluorescence of RFP and GFP, and the cell optical density (OD_600nm_). Quantitative RFP and GFP fluorescence measurements were performed via a plate reader (Infinite 200 PRO, Tecan) at an excitation wavelength of 590 nm and an emission wavelength of 650 nm for RFP and an excitation wavelength of 485 nm and an emission wavelength of 515 nm for GFP. The OD_600nm_ was determined using a spectrophotometer (Spectronic 200, Thermo scientific). Relative fluorescence intensity was calculated by normalization against OD_600nm_ of whole cells.

Autofluorescence was subtracted by measuring fluorescence of wild type strains. Fluorescence of RFP and GFP was imaged via fluorescent microscope equipped with 100× oil immersion objective lens (Revolve, Echo).

### Iron reduction measurements

Resting cell ferrozine assay^25^ was used to measure the Fe^3+^ reduction abilities of *S. oneidensis* strains. Cells from late log phase LB cultures were diluted into 5 mL fresh LB broth. Then, cells were incubated for 18 h at 30 °C under either red light or dark condition while shaking at 200 rpm. Cells were collected by centrifuging (5840R, Eppendorf) at 4,200 rpm, 4 °C for 15 min and then washed with fresh minimal medium two times. After that, cells were inoculated into sealed serum bottles containing 25 mL anaerobic minimal medium to an OD_600nm_ of about 0.1. 2 mM of ferric citrate was added into the anaerobic minimal medium as the electron acceptor. The samples were incubated at 30 °C under dark conditions without shaking. Every two hours, 10 μL of each sample was added immediately to 90 μL 1 M HCl in a 96-well plate followed by 100 μL 0.01% ferrozine. Then, after mixing the samples well and letting them sit for 10 min, the absorbance of the samples at 562 nm was determined with a plate reader (Infinite 200 PRO, Tecan). A standard curve of freshly made ferrous sulfate was used to determine the Fe^2+^ concentrations.

### Methyl orange (MO) reduction measurements

The MO decoloration assay^47^ was used to measure the MO reduction abilities of *S. oneidensis* strains. The preculture was the same as the iron reduction measurements. Cells after being cultured under either red light or dark condition were washed and inoculated into sealed serum bottles containing 25 mL anaerobic minimal medium with 100 mg/L MO as the electron acceptor to an OD_600nm_ of about 0.1. The samples were cultured at 30 °C, 200 rpm under either red light or dark condition. Absorbances of the samples from different culturing times were measured at 465 nm with a plate reader (Infinite 200 PRO, Tecan). A standard curve of absorbances of freshly made MO with different concentrations was used to determine the MO concentrations of the samples.

### Transparent-bottom bioreactor construction

Bioreactors construction was performed as our previous work^34^. Simply, planar commercial indium tin oxide (ITO) coated glass coverslips (22 mm by 40 mm) were used as the working electrodes (WEs) and the base of the bioreactors. Thin copper wires were electrically connected to the WEs with silver paint and the wire-electrode connections were then strengthened by covering with epoxy. Glass tubes (2.5 cm tall with a 20 mm and 22 mm inner and outer diameter, respectively) were adhered overtop of the WEs with siliconized sealant as the body of the bioreactors. Custom, PEEK plastic lids were used with the bioreactors along with custom Pt wire counter electrodes (CEs) and 1 M KCl Ag/AgCl reference electrodes (REs). During the cell culturing within bioreactors, the PEEK plastic lids along with CEs and REs were removed from the bioreactors and the bioreactors were simply used as culturing vessels.

### Cell culturing and biofilm formation within bioreactor

Cell culturing within the bioreactor was modified based on our previous work^34^. The late log phase LB cultures (OD_600nm_ about 1-1.5) were diluted into fresh minimal medium to an OD_600nm_ of about 0.01. 1 mL of the diluted culture was added to the bioreactor, which was made by attaching a 20 mm diameter and 2.5 cm tall glass tube on an ITO coated glass coverslip. The glass tube was sealed with a microporous membrane filter and taped to the ceiling of an incubator. A portable smart projector (A5 Pro, Wowoto) was secured below the bioreactor in the incubator and pointed up at the bottom surface of the bioreactor to shine red light with intensity of 150 μW/cm^2^. The dark samples were covered with aluminum foil to prevent undesired photoactivation. After 18 h of culturing at 30 °C in the incubator under either red light or dark condition, medium was discarded, and the biofilms on the ITO electrodes were washed three times, for 2 min each time, with minimal medium on a table shaker at 60 rpm to remove the planktonic cells. The bioreactors with fresh minimal medium were then moved to anaerobic chamber for electrochemical measurements.

### Electrochemical activity measurements

All electrochemical measurements were performed in an anaerobic chamber (Bactron 300, Sheldon Manufacturing, Inc.) with a 95:5 (N_2_/H_2_) atmosphere. Electrochemical measurements were performed with sterile minimal medium as a blank before bioreactors were used for cell culturing. After culturing and washing the biofilms, the reactor media were exchanged for anoxic media inside the anaerobic chamber before all electrochemical measurements. Biofilm cyclic voltammetry (CV) measurements were performed from -500 mV to 300 mV at 1 mV/sec using a four channel Squidstat (Admiral Instruments). Three cycles were performed for the CV and only data from the third cycle was presented in this manuscript. All potentials reported in this manuscript are vs 1 M KCl Ag/AgCl.

### Statistical analysis

All statistical analyses were performed by the Prism software (version 9.0; GraphPad) using two-tailed unpaired *t* test. All data are presented as the mean ± SD. *P* values in all graphs were generated with tests as indicated in figure legends and are represented as follows: \**p* < 0.05, \*\**p* < 0.01, \*\*\**p* < 0.001, \*\*\*\**p* < 0.0001 and ns (not significant) *p* > 0.05.

## Data availability

All relevant data supporting the key findings of this study are available within the article and Supplementary Information. Plasmids and strains generated for this study will be shared upon request to the corresponding author.

## Supporting Information

Table S1-S4: Plasmids used in this study; Strains used in this study; Primers used in this study; Recipe of *S. oneidensis* MR-1 minimal medium. Figure S1-S8: Characterization of the iLight optogenetic system without containing the heme oxygenase gene *ho1*; The thermodynamic model of the iLight optogenetic system; Expression of the iLight reporter in *S. oneidensis* strains containing different promoters of iLight repressor; Characterization of the iLight genetic circuits with different promoters of iLight repressor in *E. coli*; Characterization of the inverted iLight genetic circuits in *S. oneidensis*; Introducing of two optogenetic systems iLight and pDawn into *Shewanella*; Expression of outer membrane cytochrome MtrC through inverted iLight genetic circuit in *S. oneidensis*; Expression of periplasm cytochrome STC through inverted iLight genetic circuit in *S. oneidensis*.

## Author Information

### Corresponding Author

**James Q. Boedicker**-*Department of Physics and Astronomy, University of Southern California, Los Angeles, CA, 90089, USA; Department of Biological Sciences, University of Southern California, Los Angeles, CA, 90089, USA*

### Authors

**Fengjie Zhao**-*Department of Physics and Astronomy, University of Southern California, Los Angeles, CA, 90089, USA*

**Christina M. Niman**-*Department of Physics and Astronomy, University of Southern California, Los Angeles, CA, 90089, USA*

**Ghazaleh Ostovar**-*Department of Physics and Astronomy, University of Southern California, Los Angeles, CA, 90089, USA*

**Marko S. Chavez**-*Department of Physics and Astronomy, University of Southern California, Los Angeles, CA, 90089, USA*

**Joshua T. Atkinson**-*Department of Physics and Astronomy, University of Southern California, Los Angeles, CA, 90089, USA; Department of Civil and Environmental Engineering, Princeton University, Princeton, NJ, 08540, USA; Omenn-Darling Bioengineering Institute, Princeton University, Princeton, NJ, 08540, USA*

**Benjamin M. Bonis**-*BioTechnology Institute and Department of Plant and Microbial Biology, University of Minnesota—Twin Cities, St. Paul, MN, 55108, USA*

**Jeffrey A. Gralnick**-*BioTechnology Institute and Department of Plant and Microbial Biology, University of Minnesota—Twin Cities, St. Paul, MN, 55108, USA*

**Mohamed Y. El-Naggar**-*Department of Physics and Astronomy, University of Southern California, Los Angeles, CA, 90089, USA; Department of Biological Sciences, University of Southern California, Los Angeles, CA, 90089, USA; Department of Chemistry, University of Southern California, Los Angeles, CA, 90089, USA*

### Author Contributions

F.Z., J.T.A., J.A.G., M.Y.E.-N., and J.Q.B. designed research; F.Z. performed most of the research; C.M.N. and M.S.C. helped perform the electrochemical measurements; G.O. helped create the model; B.M.B. generated a strain used in this work; F.Z., C.M.N., G.O., and J.Q.B. analyzed data; and F.Z., G.O., and J.Q.B. wrote the paper. All authors edited the manuscript.

### Notes

The authors declare no conflict of interest.

## Acknowledgments

This study was supported by the US Office of Naval Research Multidisciplinary University Research Initiative Grant No. N00014-18-1-2632. J.T.A. was supported by the NSF Postdoctoral Research Fellowships in Biology Program under Grant No. 2010604.

## References

(1) Lindner, F.; Diepold, A. Optogenetics in Bacteria – Applications and Opportunities. FEMS Microbiology Reviews 2022, 46, 1–17.

(2) Levskaya, A.; Chevalier. A. A.; Tabor, J. J.; Simpson, Z. B.; Lavery, L. A.; Levy, M.; Davidson, E. A.; Scouras, A.; Ellington, A. D.; Marcotte, E. M.; Voigt, C. A. Engineering Escherichia coli to See Light. Nature 2005, 438, 441–442.

(3) Ohlendorf, R.; Vidavski, R. R.; Eldar, A.; Moffat, K.; Möglich, A. From Dusk till Dawn: One-Plasmid Systems for Light-Regulated Gene Expression. J. Mol. Biol. 2012, 416 (4), 534–542.

(4) Tabor, J. J.; Levskaya, A.; Voigt, C. A. Multichromatic Control of Gene Expression in Escherichia coli. J. Mol. Biol. 2011, 405 (2), 315–324.

(5) Ong, N. T.; Olson, E. J.; Tabor, J. Engineering an E. coli Near-Infrared Light Sensor. ACS Synth. Biol. 2018, 7, 240–248.

(6) Fernandez-Rodriguez, J.; Moser, F.; Song, M.; Voigt, C. A. Engineering RGB Color Vision into Escherichia coli. Nat. Chem. Biol. 2017, 13 (7), 706–708.

(7) Multam, E.; Sieryi, O.; Bykov, A.; Gerken, U.; Meglinski, I.; Andreas, M.; Takala, H. Optogenetic Control of Bacterial Expression by Red Light. ACS Synth.Biol. 2022, 11, 3354–3367.

(8) Zhao, E. M.; Zhang, Y.; Mehl, J.; Park, H.; Lalwani, M. A.; Toettcher, J. E.; Avalos, J. L. Optogenetic Regulation of Engineered Cellular Metabolism for Microbial Chemical Production. Nature 2018, 555, 683–687.

(9) Lalwani, M. A.; Ip, S. S.; Carrasco-lópez, C.; Day, C.; Zhao, E. M.; Kawabe, H.; Avalos, J. L. Optogenetic Control of the Lac Operon for Bacterial Chemical and Protein Production. Nat. Chem. Biol. 2021, 17, 71–79.

(10) Tandar, S. T.; Senoo, S.; Toya, Y.; Shimizu, H. Optogenetic Switch for Controlling the Central Metabolic Flux of Escherichia coli. Metab. Eng. 2019, 55, 68–75.

(11) Wu, P.; Chen, Y.; Liu, M.; Xiao, G.; Yuan, J. Engineering an Optogenetic CRISPRi Platform for Improved Chemical Production. ACS Synth. Biol. 2021, 10, 125–131.

(12) Jin, X.; Riedel-Kruse, I. H. Biofilm Lithography Enables High-Resolution Cell Patterning via Optogenetic Adhesin Expression. Proc. Natl. Acad. Sci. U. S. A. 2018, 115 (14), 3698–3703.

(13) Moser, F.; Tham, E.; González, L. M.; Lu, T. K.; Voigt, C. A. Light-Controlled, High-Resolution Patterning of Living Engineered Bacteria Onto Textiles, Ceramics, and Plastic. Adv. Funct. Mater. 2019, 29 (30), 1–11.

(14) Cheng, X.; Pu, L.; Fu, S.; Xia, A.; Huang, S.; Ni, L.; Xing, X.; Yang, S.; Jin, F. Engineering Gac/Rsm Signaling Cascade for Optogenetic Induction of the Pathogenicity Switch in Pseudomonas aeruginosa. ACS Synth. Biol. 2021, 10, 1520–1530.

(15) Castillo-hair, S. M.; Baerman, E. A.; Fujita, M.; Igoshin, O. A.; Tabor, J. J. Optogenetic Control of Bacillus subtilis Gene Expression. Nat. Commun. 2019, 10:3099.

(16) Pu, L.; Yang, S.; Xia, A.; Jin, F. Optogenetics Manipulation Enables Prevention of Biofilm Formation of Engineered Pseudomonas aeruginosa on Surfaces. ACS Synth. Biol. 2018, 7 (1), 200–208.

(17) Myers, C. R.; Nealson, K. H. Bacterial Manganese Reduction and Growth with Manganese Oxide as The Sole Electron Acceptor. Science 1988, 240, 1319–1321.

(18) Shi, L.; Dong, H.; Reguera, G.; Beyenal, H.; Lu, A.; Liu, J.; Yu, H. Q.; Fredrickson, J. K. Extracellular Electron Transfer Mechanisms between Microorganisms and Minerals. Nat. Rev. Microbiol. 2016, 14 (10), 651–662.

(19) McMillan, D. G. G.; Marritt, S. J.; Butt, J. N.; Jeuken, L. J. C. Menaquinone-7 Is Specific Cofactor in Tetraheme Quinol Dehydrogenase CymA. J. Biol. Chem. 2012, 287 (17), 14215–14225.

(20) Edwards, M. J.; White, G. F.; Butt, J. N.; Richardson, D. J.; Clarke, T. A. The Crystal Structure of a Biological Insulated Transmembrane Molecular Wire. Cell 2020, 181 (3), 665–673.e10.

(21) Fonseca, B. M.; Paquete, C. M.; Neto, S. E.; Pacheco, I.; Soares, C. M.; Louro, R. O. Mind the Gap: Cytochrome Interactions Reveal Electron Pathways across the Periplasm of Shewanella oneidensis MR-1. Biochem. J. 2013, 449 (1), 101–108.

(22) Edwards, M. J.; White, G. F.; Lockwood, C. W.; Lawes, M. C.; Martel, A.; Harris, G.; Scott, D. J.; Richardson, D. J.; Butt, J. N.; Clarke, T. A. Structural Modeling of an Outer Membrane Electron Conduit from a Metal-Reducing Bacterium Suggests Electron Transfer via Periplasmic Redox Partners. J. Biol. Chem. 2018, 293 (21), 8103–8112.

(23) Hartshorne, R. S.; Reardon, C. L.; Ross, D.; Nuester, J.; Clarke, T. A.; Gates, A. J.; Mills, P. C.; Fredrickson, J. K.; Zachara, J. M.; Shi, L.; Beliaev, A. S.; Marshall, M. J.; Tien, M.; Brantley, S.; Butt, J. N.; Richardson, D. J. Characterization of an Electron Conduit between Bacteria and the Extracellular Environment. Proc. Natl. Acad. Sci. U. S. A. 2009, 106 (52), 22169–22174.

(24) White, G. F.; Shi, Z.; Shi, L.; Wang, Z.; Dohnalkova, A. C.; Marshall, M. J.; Fredrickson, J. K.; Zachara, J. M.; Butt, J. N.; Richardson, D. J.; Clarke, T. A. Rapid Electron Exchange between Surface-Exposed Bacterial Cytochromes and Fe(III) Minerals. Proc. Natl. Acad. Sci. U. S. A. 2013, 110 (16), 6346–6351.

(25) West, E. A.; Jain, A.; Gralnick, J. A. Engineering a Native Inducible Expression System in Shewanella oneidensis to Control Extracellular Electron Transfer. ACS Synth. Biol. 2017, 6 (9), 1627–1634.

(26) Cao, Y.; Li, X.; Li, F.; Song, H. CRISPRi-SRNA: Transcriptional-Translational Regulation of Extracellular Electron Transfer in Shewanella oneidensis. ACS Synth. Biol. 2017, 6 (9), 1679–1690.

(27) Dundas, C. M.; Walker, D. J. F.; Keitz, B. K. Tuning Extracellular Electron Transfer by Shewanella oneidensis Using Transcriptional Logic Gates. ACS Synth. Biol. 2020, 9 (9), 2301–2315.

(28) Cao, Y.; Song, M.; Li, F.; Li, C.; Lin, X.; Chen, Y.; Chen, Y.; Xu, J.; Ding, Q.; Song, H. A Synthetic Plasmid Toolkit for Shewanella oneidensis MR-1. Front. Microbiol. 2019, 10, 1–12.

(29) Jensen, H. M.; Albersc, A. E.; Malley, K. R.; Londerd, Y. Y.; Cohen, B. E.; Helmsc, B. A.; Weigele, P.; Groves, J. T.; Ajo-Franklin, C. M. Engineering of a Synthetic Electron Conduit in Living Cells. Proc. Natl. Acad. Sci. U. S. A. 2010, 107 (45), 19213–19218.

(30) Jensen, H. M.; TerAvest, M. A.; Kokish, M. G.; Ajo-Franklin, C. M. CymA and Exogenous Flavins Improve Extracellular Electron Transfer and Couple It to Cell Growth in Mtr-Expressing Escherichia coli. ACS Synth. Biol. 2016, 5 (7), 679–688.

(31) Zacharoff, L. A.; El-Naggar, M. Y. Redox Conduction in Biofilms: From Respiration to Living Electronics. Curr. Opin. Electrochem. 2017, 4 (1), 182–189.

(32) Atkinson, J. T.; Chavez, M. S.; Niman, C. M.; El-Naggar, M. Y. Living Electronics: A Catalogue of Engineered Living Electronic Components. Microb. Biotechnol. 2023, 16 (3), 507–533.

(33) Xu, S.; Barrozo, A.; Tender, L. M.; Krylov, A. I.; El-Naggar, M. Y. Multiheme Cytochrome Mediated Redox Conduction through Shewanella oneidensis MR-1 Cells. J. Am. Chem. Soc. 2018, 140 (32), 10085–10089.

(34) Zhao, F.; Chavez, M. S.; Naughton, K. L.; Niman, C. M.; Atkinson, J. T.; Gralnick, J. A.; El-Naggar, M. Y.; Boedicker, J. Q. Light-Induced Patterning of Electroactive Bacterial Biofilms. ACS Synth. Biol. 2022, 11 (7), 2327–2338.

(35) Kaberniuk, A. A.; Baloban, M.; Monakhov, M. V.; Shcherbakova, D. M.; Verkhusha, V. V. Single-Component near-Infrared Optogenetic Systems for Gene Transcription Regulation. Nat. Commun. 2021, 12 (1), 1–12.

(36) Kreamer, N. N.; Phillips, R.; Newman, D. K.; Boedicker, J. Q. Predicting the Impact of Promoter Variability on Regulatory Outputs. Sci. Rep. 2016, 5, 18238.

(37) Berg, O. G.; von Hippel, P. H. Selection of DNA Binding Sites by Regulatory Proteins: Statistical-mechanical Theory and Application to Operators and Promoters. J. Mol. Biol. 1987, 193, 723–750.

(38) Guharajan, S.; Chhabra, S.; Parisutham, V.; Brewster, R. C. Quantifying the Regulatory Role of Individual Transcription Factors in Escherichia coli. Cell Reports 2021, 37, 109952.

(39) Kelly, J. R.; Rubin, A. J.; Davis, J. H.; Ajo-Franklin, C. M.; Cumbers, J.; Czar, M. J.; de Mora, K.; Glieberman, A. L.; Monie, D. D.; Endy, D. Measuring the Activity of BioBrick Promoters Using an In Vivo Reference Standard. Journal of Biological Engineering 2009, 3, 4.

(40) Stanton, B. C.; Nielsen, A. A. K.; Tamsir, A.; Clancy, K.; Peterson, T.; Voigt, C. A. Genomic Mining of Prokaryotic Repressors for Orthogonal Logic Gates. Nat. Chem. Biol. 2014, 10 (2), 99–105.

(41) Coursolle, D.; Gralnick, J. A. Modularity of the Mtr Respiratory Pathway of Shewanella oneidensis Strain MR-1. Mol. Microbiol. 2010, 77 (4), 995–1008.

(42) Heidelberg, J. F.; Paulsen, I. T.; Nelson, K. E.; Gaidos, E. J.; Nelson, W. C.; Read, T. D.; Eisen, J. A.; Seshadri, R.; Ward, N.; Methe, B.; Clayton, R. A.; Meyer, T.; Tsapin, A.; Scott, J.; Beanan, M.; Brinkac, L.; Daugherty, S.; DeBoy, R. T.; Dodson, R. J.; Durkin, A. S.; Haft, D. H.; Kolonay, J. F.; Madupu, R.; Peterson, J. D.; Umayam, L. A.; White, O.; Wolf, A. M.; Vamathevan, J.; Weidman, J.; Impraim, M.; Lee, K.; Berry, K.; Lee, C.; Mueller, J.; Khouri, H.; Gill, J.; Utterback, T. R.; McDonald, L. A.; Feldblyum, T. V.; Smith, H. O.; Venter, J. C.; Nealson, K. H.; Fraser, C. M. Genome Sequence of the Dissimilatory Metal Ion-Reducing Bacterium Shewanella oneidensis. Nat. Biotechnol. 2002, 20 (11), 1118–1123.

(43) Burns, J. L.; Ginn, B. R.; Bates, D. J.; Dublin, S. N.; Taylor, J. V.; Apkarian, R. P.; Amaro-Garcia, S.; Neal, A. L.; Dichristina, T. J. Outer Membrane-Associated Serine Protease Involved in Adhesion of Shewanella oneidensis to Fe(III) Oxides. Environ. Sci. Technol. 2010, 44 (1), 68–73.

(44) Gralnick, J. A.; Vali, H.; Lies, D. P.; Newman, D. K. Extracellular Respiration of Dimethyl Sulfoxide by Shewanella oneidensis Strain MR-1. Proc. Natl. Acad. Sci. U. S. A. 2006, 103 (12), 4669–4674.

(45) Luan, F.; Burgos, W. D.; Xi E L.; Zhou, Q. Bioreduction of Nitrobenzene, Natural Organic Matter, and Hematite by Shewanella putrefaciens CN32. Environ. Sci. Technol. 2010, 44 (1), 184–190.

(46) Burns, J. L.; DiChristina, T. J. Anaerobic Respiration of Elemental Sulfur and Thiosulfate by Shewanella oneidensis MR-1 Requires PsrA, a Homolog of the PhsA Gene of Salmonella enterica Serovar Typhimurium LT2. Appl. Environ. Microbiol. 2009, 75 (16), 5209–5217.

(47) Cai, P. J.; Xiao, X.; He, Y. R.; Li, W. W.; Chu, J.; Wu, C.; He, M. X.; Zhang, Z.; Sheng, G. P.; Lam, M. H. W.; Xu, F.; Yu, H. Q. Anaerobic Biodecolorization Mechanism of Methyl Orange by Shewanella oneidensis MR-1. Appl. Microbiol. Biotechnol. 2012, 93 (4), 1769–1776.

(48) Qian, Y.; Paquete, C. M.; Louro, R. O.; Ross, D. E.; Labelle, E.; Bond, D. R.; Tien, M. Mapping the Iron Binding Site(s) on the Small Tetraheme Cytochrome of Shewanella oneidensis MR-1. Biochemistry 2011, 50, 6217–6224.

(49) Ross, D. E.; Flynn, J. M.; Baron, D. B.; Gralnick, J. A.; Bond, D. R. Towards Electrosynthesis in Shewanella: Energetics of Reversing the Mtr Pathway for Reductive Metabolism. PLoS ONE 2011, 6(2): e16649.

